# Contrasting avoidance - tolerance in heat stress response from thermally contrasting climates in Arabidopsis thaliana

**DOI:** 10.1101/044461

**Authors:** Nana Zhang, Philip Carlucci, Joshua Nguyen, Jai-W Hayes-Jackson, Stephen Tonsor

**Affiliations:** Department of Biological Sciences, University of Pittsburgh, 4249 Fifth Ave, Pittsburgh, PA 15260; Carnegie Museum of Natural History, 4400 Forbes Ave., Pittsburgh, PA 15213

**Keywords:** Heat stress, avoidance, tolerance, fitness, Arabidopsis thaliana, geographic pattern

## Abstract

Plants ameliorate heat stress by avoiding heat loading, reducing tissue temperature through evaporative cooling, and/or through tolerance, i.e. maintaining function at high temperature. Here *Arabidopsis thaliana* natural populations from two ends of an elevation gradient in NE Spain were used to ask: do plants from contrasting climates 1) show genetically based differences in heat stress damage and 2) adopt different avoidance-tolerance patterns? Four low-and four high-elevation populations were repeatedly exposed to high temperature (45°C) in a growth chamber at bolting stage. High temperature induced 23% more inflorescence branches, 25% longer total reproductive branch length, and 12% less root dry mass, compared with control. However summed fruit length, hence fitness, decreased by 15%, populations did not differ significantly in fitness reduction. High elevation populations showed more avoidance, i.e. lower rosette temperature at 45°C. Low elevation populations showed more tolerance, maintaining relatively higher photosynthetic rate at 45°C. Avoidance was associated with high transpiration rate and flat rosette leaf angle. Tolerance was negatively associated with heat shock protein 101 (Hsp101) and salicylic acid (SA) accumulation. The divergent avoidance–tolerance patterns for populations from thermally contrasting climates may indicate both constraints on the evolution and contrasting adaptive divergence regulated by local climates.

## Introduction

Abiotic stresses, such as temperature and drought, are main range limitation determinants. Heat stress imposed by daily temperature fluctuation can cause severe damage to plants, including reduction of plant growth and alterations in photosynthesis and phenology. Such disruptions are ultimately likely to cause reduction in resources available for reproduction(Hasanuzzaman et al. 2013). It is therefore quite likely that frequent heat stress will reorganize allocation and physiology through selection for the highest fitness response to high temperature events. Careful observation of the relationship between a plant’s thermal environment and the specific mechanisms of adaptation to heat stress in wild populations is very limited, despite its likely relationship to extinction at the warmer end of species’ ranges.

In general, we define a heat stress as a diurnal temperature pattern in which plants display reduced fitness compared to some other temperature pattern (SØrensen 2001). Usually, the maximum stress temperature is about 10 to 15°C higher than the optimum for broadly distributed species and as little as 5°C higher than the optimum in species with narrow geographic ranges (Lindquist 1986).

Plants have developed both long-term and short-term adaptations to high temperature (Hong et al. 2003). Evolutionary adjustments of the timing of life history events (Montesinos-Navarro et al. 2011) and further adjustments in the timing of allocating of resources to rosette and inflorescence (Wolfe and Tonsor 2014), are mechanisms that allow annual plants to escape from the most stressfully high temperature period by adjusting life cycle timing. To further reduce or prevent stress from high temperature during the active growing season, adaptive responses can be described as part of two main strategies, avoidance and tolerance (Sakai and Larcher 1987).

Stress avoidance is a strategy through which plants adjust their internal states in ways that reduce exposure to a potentially damaging environment (Touchette et al. 2009, Puijalon et al. 2011). For most plants, leaves are the most important structure for obtaining energy and carbon (but see (Earley et al. 2009)). On average, avoidance can lower leaf surface temperature across growing season by 4°C compared to ambient temperature in cotton (Wiegand and Namken 1966). Generally, the higher the air temperature, the larger the differential between air and leaf can be (Linacre 1967, Wilson et al. 1987). Leaf temperature thus becomes an ideal indicator to keep track of plants’ heat avoidance.

Avoidance can also be achieved by leaf orientation adjustment (Jones and Corlett 1992, Zlatev et al. 2006). Many plants adjust leaf angle, thus reducing the leaf area that is exposed to heat from sunlight (Bradshaw 1972, Huey 2002). Populations originating from high temperature sites have higher leaf angle in heat stress in *Arabidopsis thaliana* (Vile et al. 2012) and in *Aractostaphylos* species (Shaver 1978, Ehleringer 1987, Fu 1989).

Avoidance can also be achieved through transpiration which is immediately elevated at high temperature, thus cooling the leaf surface (Shah et al. 2011). The threshold temperature that controls the relative rate of transpiration is species-specific (Mahan 1990). The transpiration process is closely connected with stomatal opening (Burke and Upchurch 1989). One potential constraint on transpiration-driven heat stress avoidance is that high temperature often co-varies with a dry environment in nature. Plants from drier and warmer sites show higher water use efficiency, compensating for large water losses due to transpirational cooling in *Boechera holboellii* populations (Knight et al. 2006). However, phylogenetic analysis of 28 dominant species in a Mexican evergreen shrubland also showed a correlation between steeper leaf angle and low transpiration rate as an adaptation to dry climate (Falster and Westoby 2003, Valiente-Banuet et al. 2010). These contrasting selection pressures on transpiration in combined heat and drought stress further complicate the evolution of avoidance in nature (Vile et al. 2012).

When internal temperatures rise sufficiently despite any avoidance mechanisms possessed by the plant, heat stress can damage cells in a variety of ways. High cellular temperature affects both cellular structural integrity and protein function, causing membrane disruption as well as disruption of metabolic function through production of reactive oxygen species (ROS) (Schöffl et al. 1998) and enzyme denaturation (Blum and Ebercon 1981, Reynolds et al. 1994, Ismail and Hall 1999). Photosynthesis is especially heat-sensitive (Berry and Bjorkman 1980, Reynolds et al. 1994, Sharkey et al. 2008). In plants measurement of extent to which photosynthetic rate is depressed can be an effective measure of functional disruption. Eventually, as temperatures rise damage is sufficient to affect a plant’s survivorship and fecundity (Senthil-Kumar et al. 2007).

Heat tolerance is the ability of plants to minimize or repair damage while experiencing a high internal temperature (Touchette et al. 2009, Puijalon et al. 2011). Heat tolerance mechanisms include protection and repair of damaged cell structures, structural proteins, and enzymes (Shah et al. 2011). While heat tolerance is complex (Kotak et al. 2007) and incompletely understood, it is known to involve up-regulation of two classes of molecules: heat shock proteins (thereafter, Hsps) (Queitsch 2000b, Hong and Vierling 2001, Wang et al. 2004) and plant hormones (Larkindale 2004, He et al. 2005, Larkindale and Huang 2005). Hsps are a group of molecular chaperones involved in dissolving and refolding aggregated cellular proteins, under both normal and stressful conditions (Hartl 1996). Members of the Hsp100/ClpB family have shown a significant role in heat tolerance in *Saccharomyces cerevisiae* (Hsp104) (Sanchez and Lindquist 1990) and *Arabidopsis thaliana* (Hsp101) (Queitsch 2000a, Hong and Vierling 2001). Plant hormones, especially salicylic acid (SA), are a common stress response. SA reduces reactive oxygen species (ROS) accumulation and affects a great many other processes in the plant. SA expression modulation and its effects are not fully understood, but SA’s importance for a variety of stress responses is well documented (Delaney et al. 1994, Klessig and Malamy 1994, Clarke et al. 2004, Yuan and Lin 2008, Vlot et al. 2009).

Plants in environments with frequent heat stress tend to evolve greater tolerance (Huey 2002), but how avoidance varies in adaptation to heat stress remains unclear. Avoidance and tolerance can have different costs and benefits depending on complex aspects of the growth environment. Thus different environments may favor tolerance or avoidance in response to the balance of selection acting on the mechanisms involved in each strategy (see for example for drought tolerance / herbivory avoidance (Siemens and Haugen 2013), for irradiance and water availability (Sánchez-Gómez et al. 2006)). It is however a mystery how avoidance and tolerance work together to contribute at the whole plant fitness level. The question of the potential relationship between avoidance and tolerance, different strategies responding to the same stimuli, is of primary ecological interest, as it may reveal constraints that limit the evolution of the traits involved in the response and variation in a broad range of architectural traits of canopies, leaves and stems, all involved in these strategies.

In this study we use a set of wild-collected *Arabidopsis thaliana* populations that exhibit clines in many traits in association with a climate gradient (Montesinos-Navarro et al. 2009, Montesinos-Navarro et al. 2011, Wolfe and Tonsor 2014). Previous work indicates that the climate gradient includes gradients in both temperature and precipitation. Thus we are particularly interested in heat stress in the context of water use.

We ask two questions. Do populations from thermally contrasting climates:

1. Display the same reductions in fitness with heat stress?
2. Exhibit contrasting avoidance and tolerance strategies?

Four low-and four high-elevation populations, each with four genotypes, were collected from northeastern Spain. In this area, climate is highly correlated with elevation (Montesinos-Navarro et al. 2009, Wolfe and Tonsor 2014). Low elevation populations experience hotter and dryer conditions, while high elevation populations experience colder and wetter conditions. In this study, we first looked at the fitness effect of repeated heat stress episodes for all the plants. We then compared avoidance and tolerance strategies in heat stress response in plants from the contrasting climates. We further explored possible causally-connected traits for both avoidance and tolerance. For avoidance, we looked at rosette angle and transpiration rate. For tolerance, we looked at the accumulation of Hsp101 and SA.

## Materials and Methods

### Materials and heat treatment

Plant lineages were collected from NE Spain as seeds (Montesinos-Navarro et al. 2009, Montesinos-Navarro et al. 2011, Montesinos-Navarro et al. 2012, Wolfe and Tonsor 2014) and grown for at least three generations in common controlled environmental conditions to remove any maternal environmental variance that might otherwise have carried over from the field. The geographicxs locations of these populations can be seen in Fig.S1 (adapted from Fig.1 in Montesinos-Navarro et al. 2001). Seeds were germinated and maintained at 22°C for 3 weeks (16 hrs light/8 hrs dark, 200 μM m^−2^s^−1^) after 5-day stratification at 5°C in the dark. Since these populations are more likely to experience heat stress at the bolting stage in nature, heat treatments were performed at bolting stage (stage 6 - 6.10 based on Table 1 in Boyes et al. 2001). Seedlings therefore experienced a 4-week vernalization at 5°C (10 hrs light / 14 hrs dark, 150 μM m^−2^s^−1^), to synchronize flowering time. After return to control growth conditions (16 hrs light/8 hrs dark, 200 μM m^−2^s^−1^) plants were checked every day and those at the bolting stage were transferred to a separate chamber for the heat treatment. Growth, control, and heat treatment were all conducted in our Conviron PGW36 controlled environment growth chambers (http://www.conviron.com) at the University of Pittsburgh. Plants from each population were blindly partitioned into two groups and randomly ordered across populations in each group. One group was the control group, in which plants were maintained at 22°C all the time, the other was the heat treatment group, in which chamber air temperature was increased steadily over 15 minutes to reach 45°C, maintained at 45°C for 3 hrs and then brought back to 22°C. This treatment was repeated twice a week from each plant’s first heat treatment till harvest (following method of Larkindale & Vierling 2008). We harvested all the plants 60 days after their first heat treatment. Avoidance and tolerance measures were performed during the first heat treatment, while fitness was estimated at harvest.

**Figure 1.**
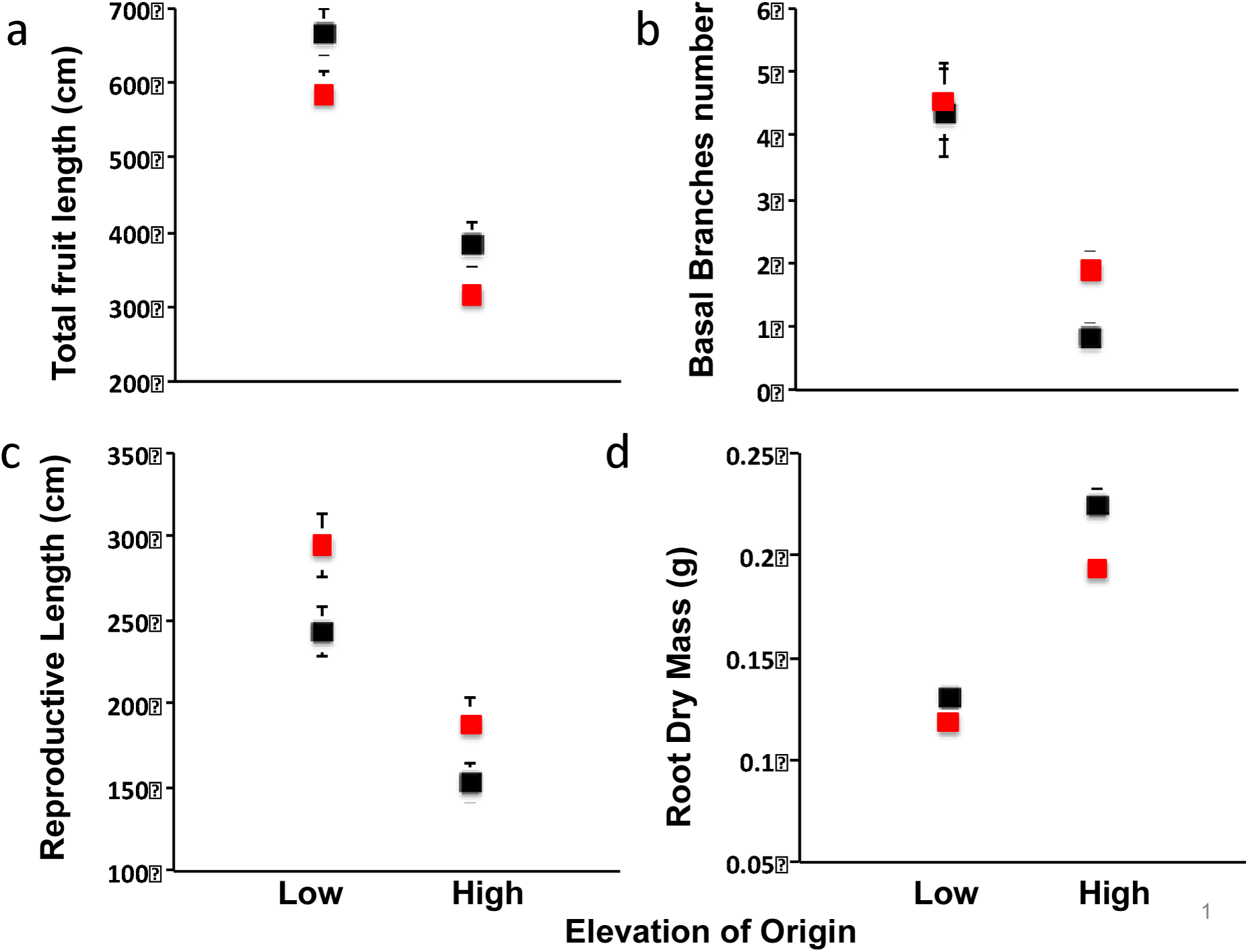
Heat stress disruption as measured by comparing trait values between 45°C heat stress and control for plants originating both low elevation and high elevations. Black = control; Red = 45°C heat stress. Figure shows the means of each elevation group under each treatment, and the bars are standard errors. See supplementary Table 1 for results from statistical analyses.

**Table 1.** ANOVA table for avoidance using Rosette temperature as a dependent variable (here DeltaT was used, which is the difference between ambient temperature and rosette temperature for normality), and rosette angle, transpiration rate, as well as heat treatment and climate of origin and their interaction as potential causal factors. This model explains 94% variation we saw in avoidance.

| Source | DF | Type III SS | Mean Square | F Value | Pr>F |
| --- | --- | --- | --- | --- | --- |
| **Full Model (R<sup>2</sup>=0.94)** | 32 | 3713.83* | 116.06 | 108.77 | <.0001 |
| Heat treatment | 1 | 1984.17 | 1984.17 | 1859.58 | <.0001 |
| Rosette Angle | 1 | 8.52 | 8.52 | 7.98 | 0.005 |
| Transpiration Rate | 1 | 0.12 | 0.12 | 0.11 | 0.74 |
| Elevation group | 1 | 0.31 | 0.31 | 0.29 | 0.59 |
| Population (Elevation group) | 6 | 236.60 | 39.43 | 36.96 | <0.0001 |
| Genotype(Population * Elevation group) | 21 | 242.67 | 11.56 | 10.83 | <0.0001 |
| Elevation group * Heat treatment | 1 | 48.69 | 48.69 | 45.63 | <0.0001 |
| **Error** | 240 | 256.08* | 1.07 |  |  |
| **Corrected Total** | 272 | 1075.98* |  |  |  |

\* Sum of Squares.

* Sum of Squares.

### Fitness quantification

We were able to measure total plant fitness 60 days after the first heat treatment, since plants were at that time nearly completely senesced. As explained in the results, there were two distinct types of fruits, aborted and mature. The distinction between the aborted and mature was visually obvious.

Sampling a substantial number of aborted fruits showed that they contained no viable seeds. Only mature fruits were included in the fitness quantification. The length of the fruit in *Arabidopsis* is highly correlated with the number of seeds within the fruit (Alonso-Blanco et al. 1999). Thus we used summed fruit length as a measure of fitness. We measured the length of five randomly chosen normal and mature fruits, two from the main (apical meristem) stem and three from secondary (lateral meristem) stems, to estimate the average fruit length. We then counted the total fruit number for each plant. Fitness, here summed fruit length, is equal to the fruit number times the average fruit length, similar to Wolfe & Tonsor (2014).

### Resource allocation quantification

We anticipated that heat stress would change resource allocation to the reproductive system and re-shape the reproductive structures, reflected in changes in reproductive branch lengths, number of branches and dry mass. To assess the potential change in resource allocation to reproduction, we partitioned the plants into rosettes, inflorescences and roots, dried them at 65°C for at least three days, and recorded the dry mass of each component. Prior to drying we also measured the length of all the reproductive portions of inflorescence branches (length from insertion of the lowest fruit to the apex) with a Map Wheel (scalex.com) and counted the number of basal branches in each plant.

### Avoidance characterization

A direct metric to indicate the level of avoidance is the difference between rosette temperature and air temperature. For rosette temperature, a thermocouple was placed in the center of the rosette, not touching the rosette surface, 15mins after the heat stress initiation. For air temperature, a thermocouple was suspended at height of the apical meristem of the tallest inflorescence in free air. For each plant the difference between rosette temperature and air temperature was calculated as the difference between the temperatures of these two thermocouples. To better understand the physiological basis for variation in rosette temperature among populations, rosette angle and transpiration rate were also quantified. To measure leaf angle, the whole rosette was photographed from four vantage points 90 degrees apart. The rosette angle is the angle of a plant’s most recently fully developed leaf to the horizontal line and was measured in ImageJ in each image (Schneider et al. 2012). We then averaged the four angles as a measure of rosette angle for each plant. We simultaneously measured transpiration and photosynthetic rate using a LiCor 6400XT gas exchange analyzer. A custom-made *Arabidopsis* single-leaf cuvette was used and one most recently fully expanded leaf was held in the cuvette until gas exchange became steady (see Fig.S2). Five measurements of carbon assimilation and transpiration were recorded at 12-second intervals and the measures were then averaged. Immediately following gas exchange measurement, each leaf was imaged and leaf area was calculated in ImageJ (Schneider et al. 2012). The photosynthetic and transpiration rates were calculated as rate per area.

### Tolerance characterization

We use relative photosynthetic rate compared with control to quantify heat tolerance. Photosynthetic rate was measured as described above with the LiCor 6400XT gas exchange analyzer. To determine the relationship of Hsp101 and SA accumulation to tolerance, we collected leaf samples and quantified Hsp101 and SA immediately following heat treatment. Two newly fully developed leaves, one for Hsp101 and the other for SA quantification, were collected right after heat treatment, freeze dried and weighed. Hsp101 quantification was measured via western blot as described in Tonsor *et al.* 2008. SA was quantified with HPLC as described in Zhang et al. 2015.

### Statistical Analyses

For each measure, we performed a separate ANOVA analysis to look at whether the measure showed significant difference at the level of elevation group, population nested within elevation group, heat treatment and the elevation group * heat treatment interaction, using Proc GLM in SAS (SAS Institute 2005). Elevation and population were treated as fixed effects.

To assess the relationship between these potential covariates and our avoidance and tolerance measures, a multivariate analysis of variance (MANOVA) was used both for avoidance and tolerance in Proc GLM (SAS Institute 2005). For avoidance, the difference between air temperature and rosette temperature, hereafter DeltaT, was treated as the dependent variable, with rosette angle, transpiration rate, heat treatment, elevation group and the elevation group * heat treatment interaction as independent variables. For tolerance, photosynthetic rate was treated as the dependent variable, while

Hsp101 accumulation, SA accumulation, heat treatment, elevation group and the elevation group * heat treatment interaction were treated as independent variables. The nested effects, population nested within elevation group, genotype nested within population, were also included in the MANOVA for both avoidance and tolerance analyses.

To further explore the direct relationship between transpiration rate and avoidance we performed univariate regression of DeltaT on transpiration rate. Likewise we tested for relationships between Hsp101/SA and tolerance by regressing photosynthetic rate on Hsp101 or SA the independent variables, using Proc REG (SAS Institute 2005).

## Results

### Heat stress caused reproductive disruption and fitness reduction

Two types of heat stress disruptions of reproduction were observed. First, in some cases heat stress damaged the apical meristem of flowering stems (Fig.S3). After the death of the apical meristem, additional secondary stems were initiated. Second, even if the apical meristem survived, heat nevertheless often led to failure of fertilization or early fruit abortion for flowers with active gametes at the time of the heat stress (Fig.S4). After heat stress, the apical meristem recovered growth but the fruits from the damaged portion of apical meristem did not successfully mature. Dissection of fruits from damaged apical meristem showed no viable seeds. We did not count such fruit in our measure of total fruit lengths.

Plants from low elevation produced greater total fruit length than plants from high elevation (p<0.0001) across both heat stress and control treatments. Across the experiment as a whole, plants from low elevation produced more basal branches (p<0.0001), greater reproductive length (p<0.0001), and lower root dry mass (p<0.0001) than plants from high elevation. Plants from both low and high elevations showed 15% reduction in total fruit length when exposed repeatedly to 45°C (p=0.0007, Fig.1a). Under heat stress, all populations showed about 25% longer reproductive length (p=0.01, Fig.1c), while only high elevation populations showed 23% more basal branches (p=0.005, Fig.1b) and 12% less root mass (p=0.0001, Fig.1d). We did not see a significant difference in the above responses to heat stress between low vs. high elevation populations (Table S1).

### High elevation populations showed greater avoidance

All plants maintained rosette temperatures that were statistically significantly different from ambient air temperature under all conditions (Fig.2). The direction of the difference in rosette temperature depended on the ambient temperature. All plants, regardless of elevation of origin, increased their rosette temperature relative to ambient temperature under the 22°C control condition. High elevation populations increased about 1.2°C more than low elevation populations (high vs. low: 24.4°C vs. 23.2°C, p<0.0001).

**Figure 2.**
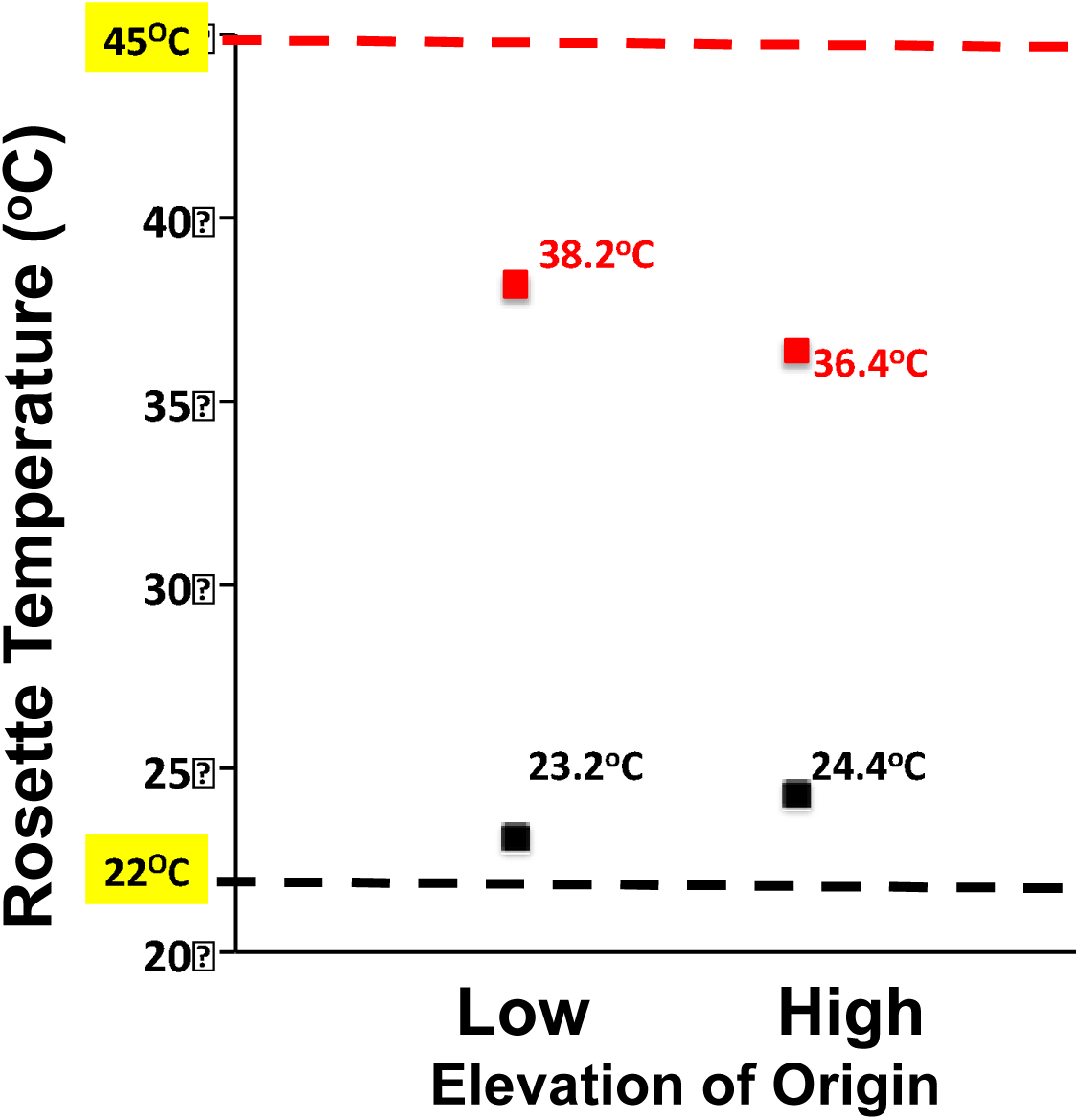
Rosette temperature, a measure of heat avoidance, comparing plants from low vs. high elevation populations, under 45°C heat stress and control. Black = control; Red = 45°C heat stress. Figure shows the means of each elevation group under each treatment, and the bars are standard errors. Dash lines show the ambient temperature for control (22°C) and heat treatment (45°C). Notice the standard error for the low vs. high elevation populations under the control is very small, so it is invisible in the figure.

When exposed to heat stress, however, both low and high elevation populations maintained rosette temperature significantly and substantially lower than ambient air temperature, on average by 7.7°C across all populations. High elevation populations reduced rosette temperature 1.8°C more than low elevation populations (low vs. high: 38.2°C vs. 36.4°C, p = 0.004). We saw greater heat stress avoidance in high elevation populations (Fig.2).

Considering both the control and heat stress treatments together, high elevation populations exhibit greater rosette temperature homeostasis than low elevation populations (Fig.2).

### Low elevation populations showed greater tolerance

Heat tolerance is the ability of a plant to perform normal plant functions when exposed to high temperature. Here we measured photosynthesis, one of the key plant functions, as a measure of tolerance (Fig. 3). Greater photosynthetic rate indicates relatively higher tolerance. We saw a significantly lower photosynthetic rate in low compared to high elevation populations under the control temperature (p<0.0001).

**Figure 3.**
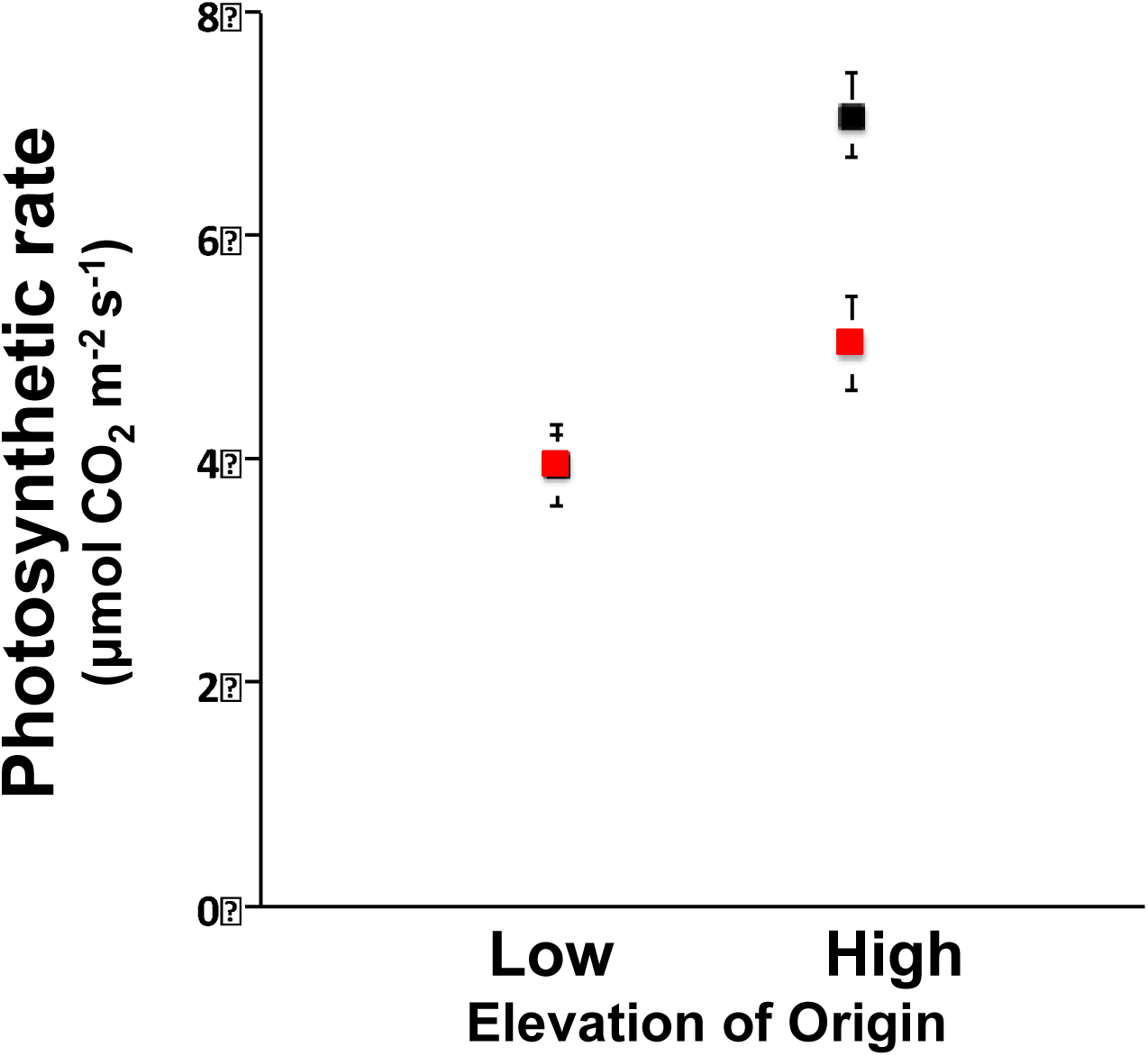
Photosynthetic rate, a measure of heat tolerance, for the low vs. high elevation populations under the 45°C heat stress and control. Black = control; Red = 45°C heat stress. Figure shows the means of each elevation group under each treatment, and the bars are standard errors. Note the photosynthetic rates for low elevation control and 45°C heat stress treatments overlap each other.

However, with a 45°C heat stress, low elevation populations showed no significant change in photosynthetic rate, while high elevation populations significantly reduced their photosynthetic rate (Elevation group*Heat treatment interaction: p=0.003, Table S1). Low elevation populations were significantly more heat tolerant than high elevation populations (p<0.0001).

### Avoidance was positively associated with high transpiration rate and flat rosette angle

We measured rosette angle and transpiration rate as potential traits associated with avoidance (Fig. 4, Fig.5, Table S1).

**Figure 4.**
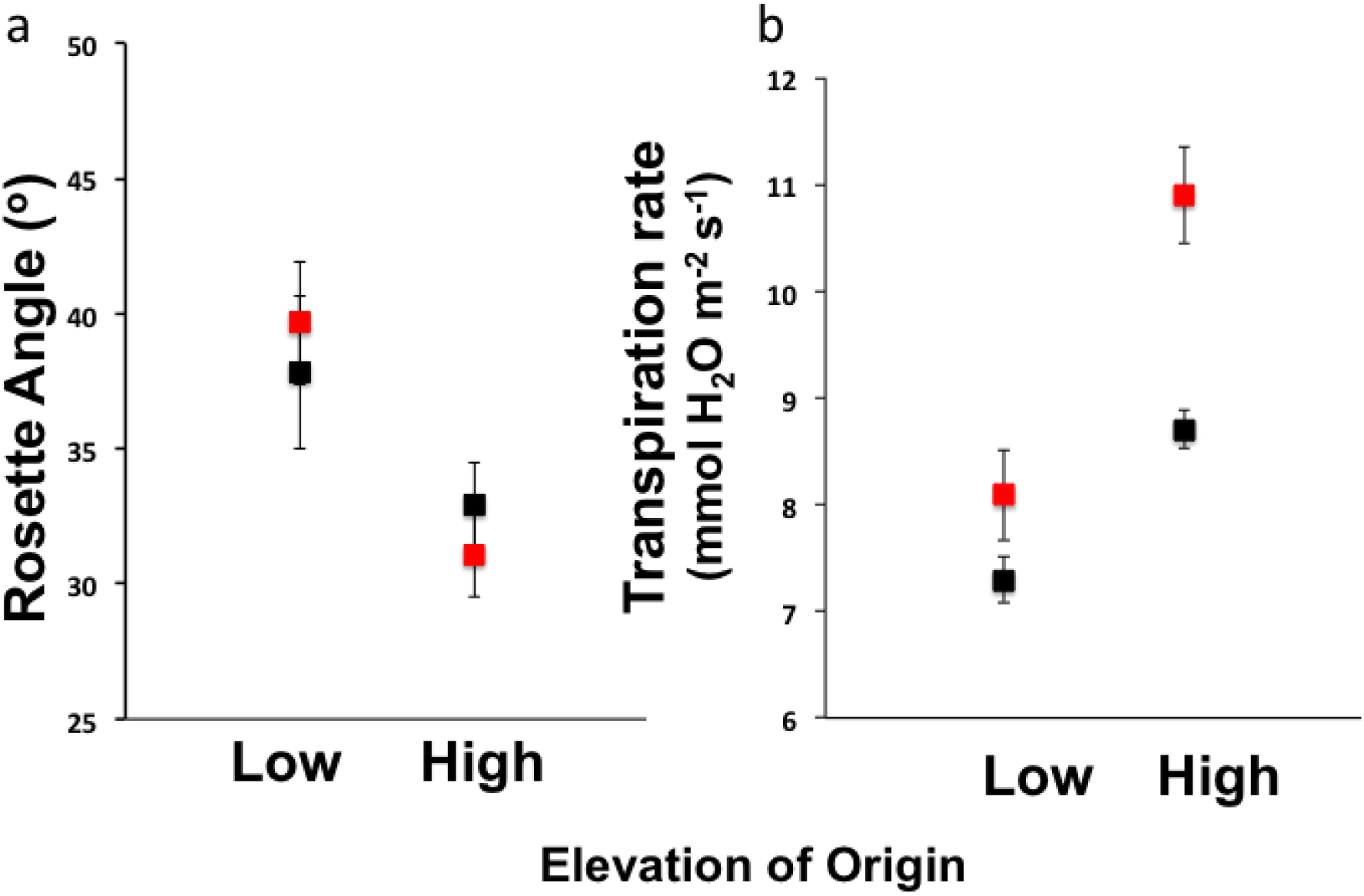
Two potential avoidance mechanisms compared for the low vs. high elevation populations under 45°C heat stress and control. Black = control; Red = 45°C heat stress. Figure shows the means of each elevation group under each treatment, and the bars are standard errors.

**Figure 5.**
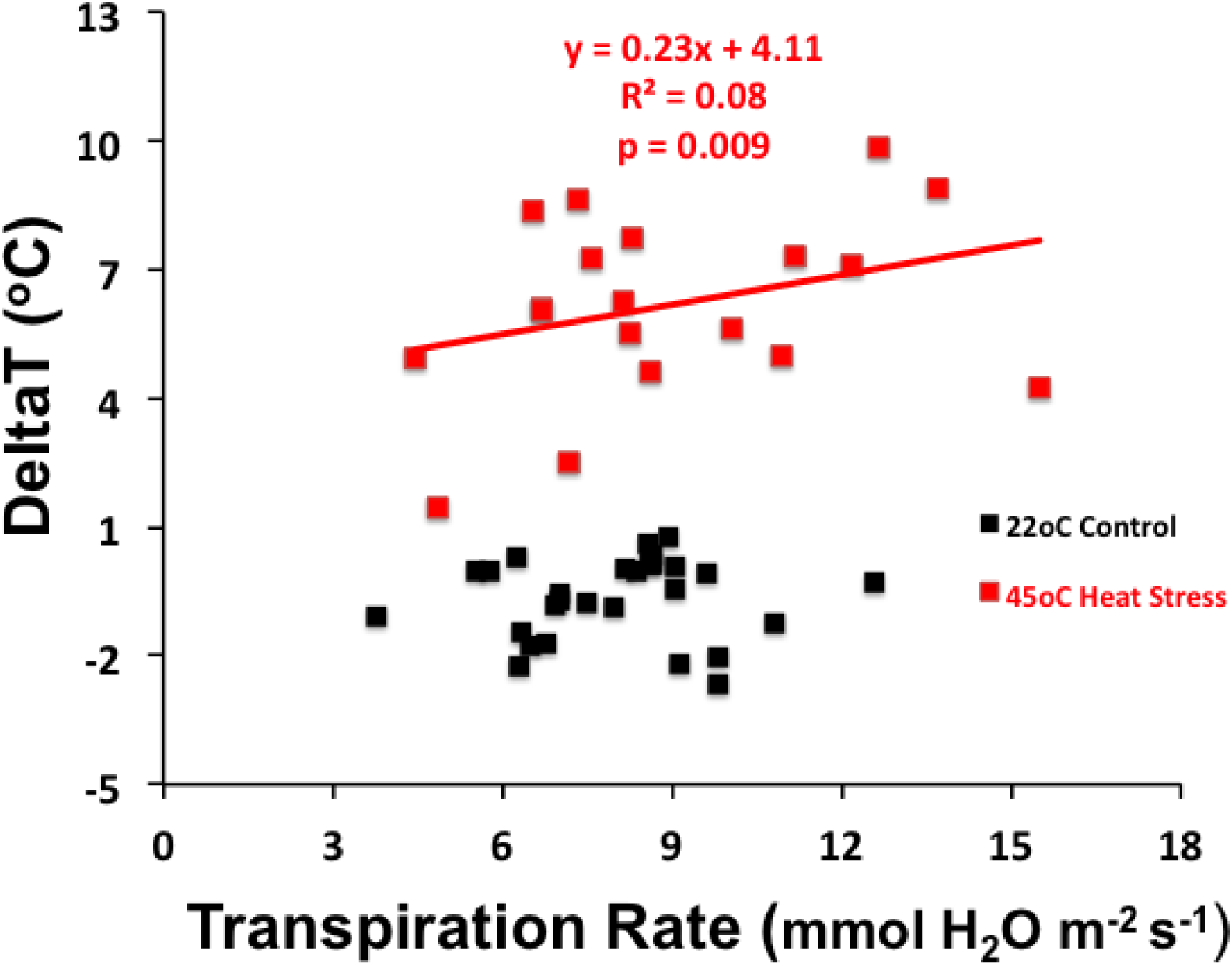
The relationship between transpiration rate and DeltaT (the difference between air and rosette temperature), comparing 45°C heat stress and control. Data points displayed are genotype means at both control and heat treatment groups. Statistical analysis was done with individual plant trait values (total samples = 84) using Proc REG in SAS. The regression line for the 45°C heat stress is: DeltaT = **0.23\*** Transpiration rate **+ 4.11**. The slope is statistically significant (p=0.009).

Rosette angle differed between low and high elevation populations regardless of treatment, with low elevation populations exhibiting sharper rosette angle (low vs. high mean rosette angle: p=0.0005). However, under our measurement protocol the rosette angle was not significantly affected by heat treatment (Fig. 4a).

High elevation populations showed significantly higher transpiration rate than low elevation populations under the control temperature (p<0.0001, Fig.4b). Transpiration rate was significantly increased with heat treatment (p<0.0001), with high elevation populations increasing significantly more than low elevation populations (p=0.0002).

Our MANOVA analysis explained 94% of the variation in avoidance (p<0.0001, Table 1). Significant interaction effects were observed between elevation groups and heat treatment (p<0.0001), indicating that low and high elevation populations have evolved different responses to heat. We also saw a significant effect of rosette angle (p=0.005). However, we did not see a significant effect of transpiration rate. We observed highly significant nested effects for both population and genotype (p<0.0001 for both).

To further explore the direct relationship between transpiration rate and avoidance, separate regression analyses in the two heat treatments of DeltaT on transpiration rate showed a significant positive relationship between transpiration rate and DeltaT at 45 °C (p=0.009, Fig.5), e.g., higher transpiration rate was associated with higher DeltaT. We did not see a significant relationship between transpiration rate and DeltaT in the control (Fig.5). Even though we did not detect a significant effect of transpiration rate on DeltaT in the MANOVA analysis, this separate analysis confirmed its direct effect on DeltaT at 45 °C. We consider these apparently contradictory results for transpiration rate’s effects in the discussion.

### Tolerance was negatively associated with Hsp101 and SA accumulation

With heat stress, Hsp101 accumulation was significantly increased 22 fold and 8 fold for low and high elevations populations, respectively (Fig. 6a, p<0.0001, Table S1). However, because of the large variation within each elevation group, we did not detect a significant difference between low vs. high elevation populations in Hsp101 accumulation in the heat treatment (Fig.6a).

**Figure 6.**
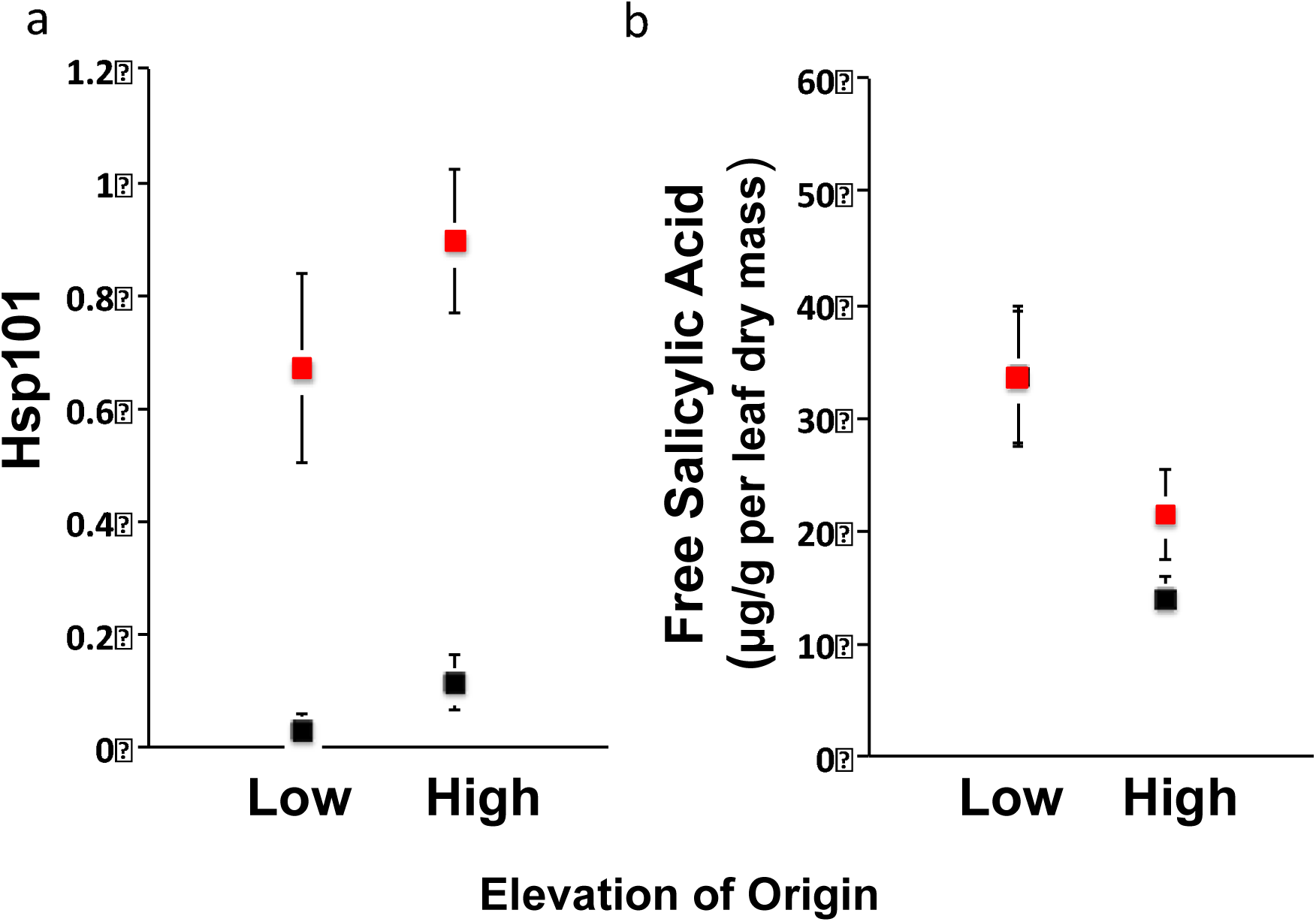
Two tolerance mechanisms, Hsp101 ratio and free salicylic acid, and their relative value in low vs. high elevation populations under 45°C heat stress and control. Black = control; Red = 45°C heat stress. Figure shows the means of each elevation group under each treatment, and the bars are standard error. Notice in b) the free salicylic acid concentration for the control overlapped with its concentration at the 45°C heat stress. The Hsp101 ratio is its concentration relative to our biological standard, in which we heat treated a combination of leaf samples at 45°C and used it as a quality control on gel-to-gel variation in western blot, see details on materials and methods in Tonsor et al. 2008. The unit for b) free salicylic acid in the figure is ug/g per leaf dry mass.

Both free and total SA were higher in low elevation populations than in high elevation populations across the experiment as a whole (Fig. 6b, free salicylic acid: p=0.0001; total salicylic acid data not shown). However, with heat treatment, high elevation populations significantly increased free and total SA, while low elevation populations showed no significant difference compared to control (Fig. 6b) in the 45°C heat stress.

Our MANOVA explained 93% of the variation in tolerance (p<0.0001, Table 2). Significant effects of Hsp101 accumulation (p=0.005), total SA (p<0.0001), free SA (p=0.005) were observed. The interaction between elevation group and heat treatment (p=0.01) was also significant (Table 2), indicating evolved differences in elevation groups in their response to high temperature.

**Table 2.** ANOVA table for tolerance using photosynthesis rate as a dependent, and Hsp101, free and total salicylic acid as well as heat treatment and climate of origin and their interaction as potential causal factors. This model explains 93% variation we saw in tolerance.

| Source | DF | Type III<br>SS | Mean<br>Square | F Value | Pr>F |
| --- | --- | --- | --- | --- | --- |
| **Full Model (R<sup>2</sup>=0.93)** | 32 | 1384.38* | 43.26 | 47.36 | <0.0001 |
| Heat treatment | 1 | 1.95 | 1.95 | 2.13 | 0.14 |
| Elevation group | 1 | 64.53 | 64.53 | 70.63 | <0.0001 |
| Population (Elevation group) | 6 | 243.19 | 40.53 | 44.37 | <0.0001 |
| Genotype(Population * Elevation group) | 20 | 617.81 | 30.89 | 33.81 | <0.0001 |
| Hsp101 accumulation (log) | 1 | 7.41 | 7.41 | 8.11 | 0.005 |
| Total salicylic acid accumulation (log) | 1 | 73.11 | 73.11 | 80.02 | <0.0001 |
| Free salicylic acid accumulation (log) | 1 | 7.51 | 7.51 | 8.22 | 0.005 |
| Elevation group * Heat treatment | 1 | 5.90 | 5.90 | 6.46 | 0.01 |
| **Error** | 112 | 102.32* | 0.91 |  |  |
| **Corrected Total** | 144 | 1486.70* |  |  |  |

\* Sum of Squares.

* Sum of Squares.

Univariate regression analysis of photosynthetic rate on Hsp101 accumulation showed a significant negative relationship, e.g., higher Hsp101 accumulation was associated with lower photosynthetic rate (p=0.03, data not shown). A negative relationship was also found between photosynthetic rate and free SA accumulation (p=0.005). These two univariate analyses are concordant with the results of the MANOVA.

## Discussion

Here we have shown that a 45°C repeated heat stress imposed periodically starting at the bolting stage is a significant heat stress for genetic lines collected from natural populations of *Arabidopsis thaliana* in NE Spain, since it caused significant decrease in fruit production compared to a benign control temperature (Fig.1). We then showed that, although both avoidance and tolerance were observable in all populations in response to heat stress, high elevation populations manifested more avoidance (Fig.2) and low elevation populations showed more tolerance (Fig.3). Our mechanistic analyses further showed that avoidance was positively associated with high transpiration rate and flat rosette angle (Fig.4, Fig.5, Table 1), while tolerance was negatively associated with Hsp101 and SA accumulation (Fig.6, Table 2). The 8 populations used in this study are part of 17 populations along a climate gradient described in previous studies. In those prior studies we observed strong clines in many traits (Montesinos-Navarro et al. 2009, Montesinos-Navarro et al. 2011, Montesinos-Navarro et al. 2012, Wolfe and Tonsor 2014). We previously showed that population genetic analyses strongly support the hypothesis that this cline results from local adaptation along a climate gradient associated with altitude (Montesinos et al. 2009, Montesinos-Navarro et al. 2011). Likewise the contrast in strategy in low vs. high elevation populations observed in this study indicates differential evolutionary responses to heat stress associated with adaptation to Mediterranean low elevation vs. interior high elevation climates.

Despite the importance of heat stress, there is very little work that examines genetically based adaptive differentiation among lineages in heat avoidance and heat tolerance. However, much work has been done on other abiotic stresses. Abiotic stresses, drought stress and salt stress in particular, show a similar pattern of avoidance vs. tolerance in various plant species. Farrant et al. (1999) reported a negative relationship between drought avoidance and tolerance in three desiccation-tolerant angiosperm species (M.Farrant et al. 1999). Five herbaceous wetland plant species showed varying combinations of avoidance and tolerance in response to short-term drought stress (Touchette et al. 2007). Four of the species showed an avoidance strategy while all five species also showed a tolerance strategy. Rahman et al. (2011) compared the relative contribution of avoidance and tolerance to drought stress in two kiwifruit species, finding *Actinidia deliciosa* had lower avoidance and higher tolerance than *Actinidia chinensis* (Rahman et al. 2011). Similarly, Touchette et al. (2009) showed a contrasting response to salt stress in marsh halophytes *Juncus roemerianus* and *Spartina alterniflora*, in which *Juncus roemerianus*, experiencing transient salt stress exposure, showed salt avoidance and *Spartina alterniflora*, with frequent long-term salt exposure, showed salt tolerance (Touchette et al. 2009). A diverse array of abiotic stresses share some common pathways at both physiological and molecular levels (Pastori and Foyer 2002), suggesting that we might expect a similar pattern as we learn more from heat stress. Further exploration of the constraints on the evolution of the two strategies at morphological, physiological, genomic and gene expression level can provide insights in understanding the distribution pattern of plants and how adaptive responses evolve over time.

The contrast in evolved relative importance of avoidance and tolerance between our two climatic study regions indicates disruptive selection on heat stress response between the high and low elevation regions of our source populations. This disruptive selection, selecting for greater avoidance at high elevation but greater tolerance at low elevation, is likely the result of different costs for each strategy depending on the local physical environment.

In *Impatiens capensis* early season drought stress selects for avoidance but later drought stress favors tolerance (Heschel & Riginos 2005). *Nicotiana tabacum* shows a sequential response in drought stress, first avoidance then tolerance, indicating avoidance is favored in short-term stress but tolerance is favored in long-term stress (Riga and Vartanian 1999). This is in accordance with our study in *Arabidopsis thaliana* as well. When the observed avoidance and tolerance patterns were put in the context of climate, we saw interesting associations of response to heat with climate of population origin. For example, we know that annual precipitation in high elevation is 550mm greater than precipitation in low elevation (Wolfe and Tonsor 2014) and continues longer into the summer season (Montesinos-Navarro et al. 2009). This may allow greater transpiration in high elevation populations, contributing to the greater ability to avoid high temperature we observed among high elevation populations. Studies on drought and salt stress also showed that plants that have greater access to water adopt avoidance rather than tolerance (Touchette et al. 2007, Touchette et al. 2009). Similarly, the average annual temperature is up to 11°C higher in our low elevation sites compared to high elevation sites (Wolfe and Tonsor 2014); thus populations from low elevation are constantly exposed to higher temperatures compared to high elevation populations. This, combined with the lower availability of water for transpirative cooling, may explain why low elevation populations are more tolerant and less resistant than high elevation populations.

We found an increase in rosette temperature in the 22°C control but a decrease in the 45°C heat treatment compared to the ambient temperature for all populations. The difference between rosette temperature and air temperature is positive in cool but negative in hot air. The air temperature at which one observes zero air-leaf temperature differential has been called the “equality temperature” (Linacre 1964). The equality temperature is often around 30°C in well-watered, thin-leaved plants (Linacre 1967) but is species-specific (Savvides et al. 2013). For example, cotton has an equality temperature of 27°C (Upchurch and Mahan 1988). Based on our data and assuming a linear response, we can draw approximate equality temperatures at 25.4°C and 27°C for low elevation and high elevation populations, respectively. This 1.6°C difference in equality temperature reflects an intraspecific differentiation in homeostatic control in natural *Arabidopsis* populations. In a range of 22 – 45°C, high elevation populations were more homeostatic than low elevation populations (Fig.2). Our study also supported Mahan and Upchurch’s (1988) proposal that plants are capable of at least limited homeothermy.

Transpirational cooling is one of the most important transient avoidance mechanisms in plants (Burke and Upchurch 1989). The importance of transpiration and homeostatic control in meristem temperature has been shown for cucumber and tomato plants (Savvides et al. 2013), as well as cotton (Burke and Upchurch 1989). Our study also revealed a direct positive relationship between the transpiration rate and avoidance (Fig.5), even though we could not detect a significant effect in the combined MANOVA analysis (Table 1). Measures of transpiration rate are noisy, especially at high temperature. *Arabidopsis* populations exhibit a high level of genetic homogeneity. Most of the measured trait variance in studies of natural populations of *Arabidopsis* is between populations and regions (e.g. Montesinos et al. 2009). The certainty of assignment to population and elevation, the high trait variance among populations and regions, and the high error variance in transpiration measures mean that most of the causal variance is absorbed at the population and region level in the MANOVA, leaving little variance directly attributed to transpiration rate.

It is also important to emphasize that our transpiration rate measures were conducted on a single leaf. Transpirational cooling of the rosette involves interactions of the complex stacked leaf structure of the rosette and its interaction with the micro-climate in which plants reside. The rosette temperature depends in a complex way on the aggregate functioning of all the individual leaves in the rosette. Whole rosette transpiration is influenced not only by the leaf properties under uniform conditions, but also on the much more complex influence of rosette structure. The structure leads to variation in the rates of energy loading to individual leaves. It also determines convective and conductive transfer of heat to the surrounding air. These factors further influence the steepness of the water vapor diffusion gradient, and temperature gradient around individual leaves. A full understanding of control over air and tissue temperature within rosettes will require study of the rosette as a functional unit.

Leaf angle did not show the significant hyponastic response we expected in this study. This is likely because we measured this trait too soon after heat stress. A constitutively steep leaf angle is a long-term adaptive trait to deal with water deficit, high radiation load, or high temperature (Fu 1989, Falster and Westoby 2003, Valiente-Banuet et al. 2010). Plants from low latitude showed much steeper leaf angle compared with high latitude plants in 21 European *Arabidopsis thaliana* ecotypes, and all plants displayed steeper leaf angle in response to extended vernalization period (10, 20, or 30 days) (Hopkins et al. 2008). Physiologically-driven changes in leaf angle can be elicited by a variety of environmental conditions, including heat. Leaf angle movement generally requires observation over a number of hours (Ehleringer 1987). However, in our study, we measured leaf angle 15 min after heat treatment started. We hypothesize the leaf angle might change if given repeated and prolonged heat stress period and measured later in the treatment.

Hsp101 and SA accumulation both had very high variation (Fig.6) in this study. In prior published studies plants were assayed for Hsp101 and SA accumulation in seedling stage when plants have not yet developed any functional avoidance mechanisms. Seedlings therefore experience a heat stress temperature that is equal to the ambient heat treatment temperature. Thus in these prior studies, Hsp101 and SA accumulation is less variable and is distinguishable among populations (Tonsor et al. 2008, Zhang et al. 2014, Zhang et al. 2015a). However, adult plants can adopt both avoidance and tolerance in heat stress response. As shown in Fig.2, the actual rosette temperature for a 45°C heat stress is 36.4°C for high elevation populations vs. 38.2°C for low elevation populations, on average. Hsp101 accumulation increases rapidly in the range of 34-40°C among *Arabidopsis* plants collected from natural populations (Tonsor et al. 2008). The observed variation in this study in the actual rosette temperature might explain why we did not detect a significant difference in Hsp101 accumulation between low vs. high elevation populations at the 45°C heat stress (Fig.6a); based on our past studies the various genetic lines used here are very likely to be variable in expression in uniform temperature, and differ in their DeltaT. As a result they are highly variable in their responses in an experiment like the one reported here. To the best of our knowledge this is the first time that Hsp101 has been quantified in adult plants that experienced variable plant tissue temperature despite uniform ambient temperature. In addition our Hsp101 measurement method and Hsp101 expression itself are both highly variable at the level of both biological and technical replicates (data not shown). In this study, we estimated the necessary sample sizes based on prior seedling experiments, under conditions in which variation in heat avoidance was not possible. However, in retrospect based on the results of this study, an estimated sample size of 96 (48 samples for each elevation group) would be needed to distinguish the difference in low and high elevation populations in their response to the 45°C heat.

In a previous experiment, we detected a cline in SA in genetic lines collected along our study system’s elevation gradient, when measured in a 22°C environment (Zhang et al. 2015b). In the present study, even with the large variation in rosette temperature observed, we still detected a significant difference in free SA accumulation comparing low and high elevation populations at both control and 45°C heat stress temperatures (Fig.6b). The total SA accumulation for the low elevation populations was indistinguishable in the control vs. the 45°C heat treatment, while the total SA value was significantly increased by about 180% in the 45°C heat treatment for the high elevation populations, compared with their control (data not shown

Hsp101 and SA expression are both rapidly up-regulated with heat stress. We observed a negative association between both Hsp101 and SA and photosynthetic rate. This suggests that the accumulation of Hsp101 and SA are upregulated when cellular or subcellular damage is sensed by the plant, i.e. the same conditions in which photosynthesis declines.

The high avoidance ability under high temperature in adult plants, as we saw in Fig.2, indicates the necessity of connecting lab studies with more accurate reflections of field conditions. Previous heat stress response studies focus on seedlings under controlled lab conditions, yet it is at the reproductive stage that Arabidopsis and other spring annuals and biennials most often encounter high temperatures.

Because our study and others (Helliker and Richter 2008, Broitman et al. 2009, Helmuth et al. 2010) demonstrate that adult plants can maintain leaf and rosette temperature that differs substantially from ambient temperature due to the avoidance mechanisms, studies of heat tolerance and stress responses will be most fruitful if done in reference to plant tissue temperature instead of ambient temperature. However, even with these refined and comprehensive measures, mysteries still exist regarding how whole plants respond to heat stress in nature. Transcriptome sequencing data from these low vs. high elevation populations under heat stress could provide more detailed information about the gene networks for universal and regional-specific stress response.

## Contributions by authors

NZ and SJT designed the experiment. NZ, PC, JN and JH performed the experimental work. NZ and SJT performed analyses. All co-authors discussed and interpreted results prior to writing. Writing was done by NZ and SJT.

## Acknowledgement

We are grateful to Tonsor Lab managers Kali Theis and Tim Park for excellent technical assistance. Elizabeth Vierling, University of Massachusetts, Amherst, generously provided Hsp101 antibody and advice. Jeff Brodsky, University of Pittsburgh, was generous in logistical support and advice for western blot assays of Hsp101. SJT is extremely grateful to F. Xavier Picó of Estación Biológica de Doñana, Seville, Spain for introduction to the Spanish Arabidopsis system and for many hours of friendship during field characterization and collection of the populations.

---

**Figure S1.** Geographic location of the 8 populations (adapted from Fig.1 in Wolfe and Tonsor 2014, New Phytologist). Map showed all 16 Arabidopsis thaliana populations in northeastern Spain. The eight populations used in this study were highlighted with black dots.

**Figure S2.** Using single leaf cuvette to quantify photosynthetic rate and transpiration rate. Plants shown were at the pre-bolting stage. Here we only use this photo to show the equipment used for quantification. We do not have a figure to show the measurement at the bolting stage.

**Figure S3.** Heat stress caused more inflorescences to be induced. Red arrows showed the regions where the stems were damaged, and green arrows showed new branches initiated after heat stress. Figure shows both the whole reproductive part and the detailed heat stress disruption of shoot apices.

**Figure S4.** Heat stress reduced fruit quality. Damage occurred in the middle of the main stem at a time when that stem segment was near the stem apex. After heat stress, the main stem growth recovered, but the fruits did not succeed to mature, possibly because of pollen inviability. Dissection on fruits of this kind showed that they did not produce viable seeds. Figure shows both the damage on the whole reproductive part and a enlarged view of section of the stem containing heat-damaged fruit. In the enlargment, the short red lines mark damaged fruits and the short green lines mark normally developed fruit. The damaged fruits are much smaller compared to a normal fruit. The smaller dashed rectangles highlight additional stems with similar damage.

**Table S1.** ANOVA table for each variable measured in this study.

